# Rapid and reproducible *in vitro* generation of human parvalbumin-expressing cortical interneurons

**DOI:** 10.64898/2026.03.09.710579

**Authors:** Karima Azzouni, Daniel D’Andrea, Asmaa Ghazwani, Sophia Wilson, Andrew J. Pocklington, Eunju Shin

## Abstract

Parvalbumin-expressing cortical interneurons play a critical role in maintaining the balance between excitatory and inhibitory signalling and are essential for cognition, with dysfunction implicated in numerous brain disorders. Although human pluripotent stem cells have enabled the generation of diverse human neuronal types *in vitro*, including cortical interneurons, parvalbumin-expressing interneurons - unlike somatostatin-expressing interneurons - remain difficult to generate reliably and consistently. Here, we demonstrate the efficient and reproducible generation of parvalbumin-expressing cortical interneurons *in vitro* within 50 days of differentiation. Parvalbumin mRNA and protein were detected without forced gene expression, cell sorting, rodent co-culture or intracerebral transplantation, approaches commonly required by previous protocols. Single-cell transcriptomic analyses validated neuronal identity and authenticity, revealing enrichment for gene expression signatures of parvalbumin-expressing cortical interneurons *in vivo*. Together, these findings establish a robust method that facilitates interneuron research by enabling the reliable generation of authentic human parvalbumin-expressing cortical interneurons within a short time frame.

**eTOC blurb:** Azzouni *et al*. present a rapid and reproducible protocol for generating authentic human parvalbumin-expressing cortical interneurons from pluripotent stem cells in just 50 days, without forced gene expression or co-culture. Single-cell transcriptomics confirm robust acquisition of *in vivo*-like PVALB interneuron identity, enabling new opportunities for human interneuron research.

**Highlight:**

- Optimising SHH and WNT modulators enables consistent PVALB interneuron generation.
- 10% of cells express PVALB mRNA within 50 days of 2D differentiation from hPSCs.
- PVALB expression occurs without gene forcing, sorting, co-culture or grafting.
- Comparison of gene expression to *in vivo* interneurons confirms PVALB authenticity.

**Graphical abstract:** 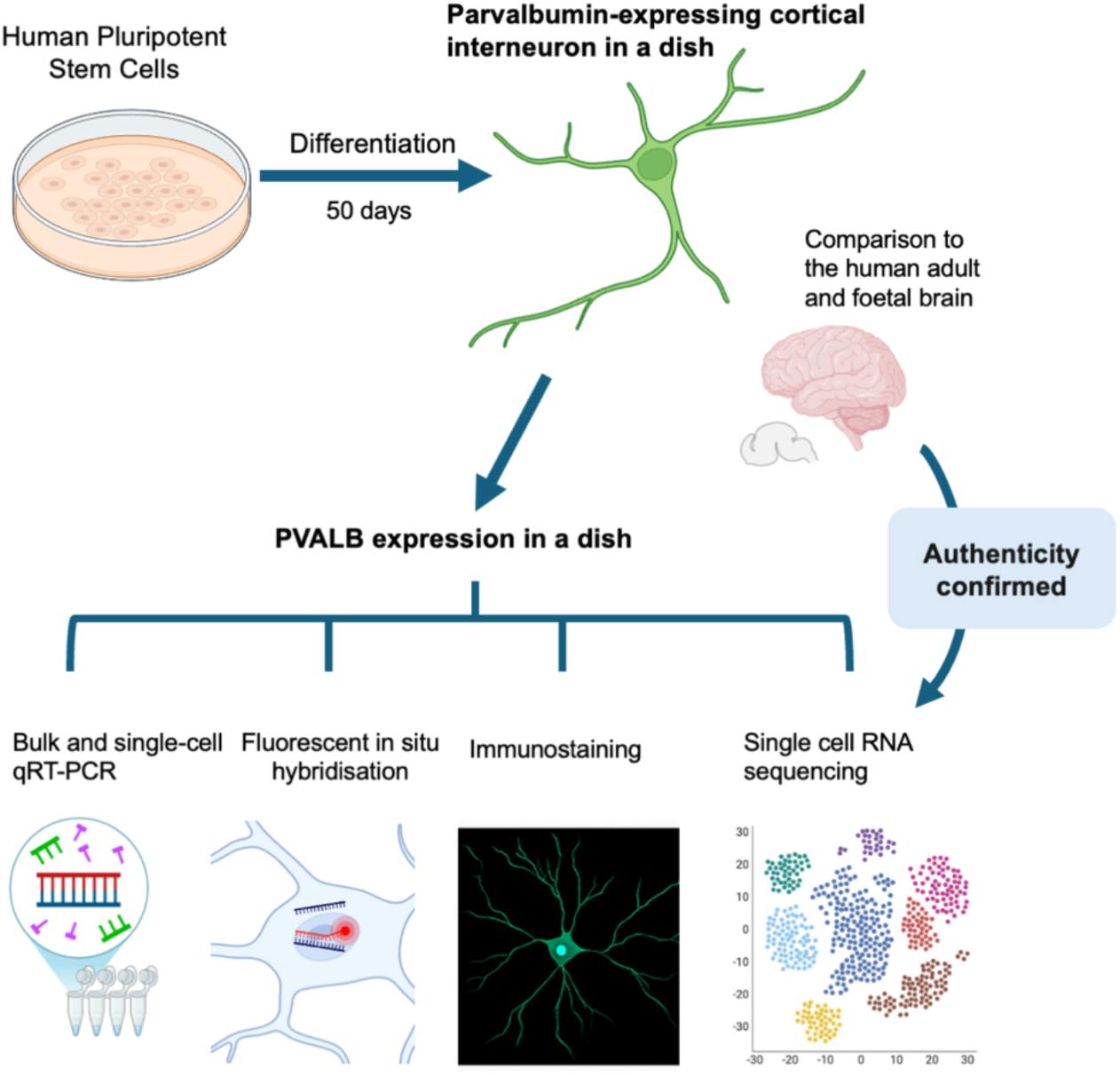

## Introduction

Cortical interneurons (CIs) play a pivotal role in balancing out the excitatory signals produced by cortical projection neurons and are postulated to be involved in several psychiatric and neurological disorders^1,2^. They are generated from the ventral forebrain, with various signalling molecules interacting to form their identity. There are three main types of cortical interneurons: Parvalbumin, Somatostatin and 5HT_3a_R expressing interneurons^3^. The first two are generated from medial ganglionic eminence (MGE) while the last type is from caudal ganglionic eminence (CGE), which can also be further divided into sub-types such as VIP, LAMP5 and PAX6 expressing neurons^4^. These different types of CIs input into different locations within pyramidal neurons in the cortex, regulating distinct pathways and functions^3^.

Much of this knowledge has been gained through research using mouse brains. Due to significant human-rodent differences in cortical organisation and function, including cognitive ability, investigation of human neurons is necessary. Since access to human neurons is limited and impractical, the CIs used in most disease modelling research are derived from human pluripotent stem cells (hPSCs). There are several published protocols for generating MGE-derived interneurons^5–7^ and CGE-derived interneurons^8^. While MGE-derived SST and CGE-derived VIP neurons are frequently and stably generated by these protocols, MGE-derived PVALB interneurons have been challenging to produce: no study has yet been able to demonstrate the production of PVALB-expressing interneurons at levels detectable via scRNAseq in 2D culture. This could reflect the natural developmental timeline of PVALB interneurons and/or suboptimal culture conditions, which seek to recreate the local micro-environment within the area of the MGE where PVALB neurons are born. Most protocols employ activation of SHH and inhibition of WNT signalling^5,6^, as these are the main patterning factors in the MGE. Some also add FGF8 to rostralise cells and counteract the caudalising effect of FGF19 caused by SHH activation^7^. While the timing and concentration of extrinsic factors required to generate MGE-derived cortical interneurons have been investigated, studies to date have not specifically investigated the optimum dose of extrinsic factors required to generate PVALB interneurons. As PVALB and SST interneurons are preferentially born in ventral and dorsal MGE, respectively^9^, and SHH concentration is higher dorsally than ventrally in MGE^10^, we postulated that different optimum concentrations of extrinsic factors such as SHH would be required to produce either PVALB- or SST-enriched interneuron populations. In addition, while the effect of varying individual extrinsic factors has been investigated, a combinatorial approach in which the concentrations of multiple factors are simultaneously varied has not. We reasoned that a combinatorial approach would be more likely to capture any subtle interactions between extrinsic factors required to produce cortical interneurons *in vitro*.

In this study, we tested 12 different combinations of SHH and WNT inhibitor XAV939 to find the best culture conditions to generate PVALB alongside other MGE interneurons. Four conditions were selected for further testing in three different hPSC lines, where single-cell analysis - including single-cell qRT-PCR and single-cell RNA sequencing - was performed to validate the identity of PVALB cortical interneurons generated through our protocol.

## Results

### Comparison of 12 combinations of SHH and XAV939 concentrations reveals one optimal condition that favours PVALB expression

Our initial differentiation protocol was devised based on several published papers^5–7^. We were able to confirm the rostralising effect of FGF8 noted by Kim et al., 2014^7^, and thus included FGF8 in our culture (Figure S1). While the treatment window and concentration of SHH and XAV939 have been separately tested in previous studies, with the aim to generate MGE-derived cortical interneurons (those from NKX2.1^+^ neural precursors), we are not aware of any study investigating combinations of these molecules in different concentrations. In addition, none of the studies to date have aimed to find the optimal concentration of these factors to generate PVALB interneurons. Given the potential for interaction between these factors - both of which are crucial for patterning - we tested three XAV939 concentrations (0, 1 and 2µM) and four of SHH concentrations (0, 50, 100 and 200ng/ml) in combination, totalling 12 conditions (Figure 1A) to find the optimal condition to generate PVALB cortical interneurons. Other patterning molecules that were included in all the conditions are LDN193189 100nM and SB43152 10µM to make cells into neuroectoderm, and the SHH agonist Purmorphamine 1µM and FGF8 100ng/ml. Purmorphamine is a small molecule SHH agonist, so cellular SHH signalling was activated in all conditions: the different SHH concentrations allowed us to fine-tune this signalling. FGF8 was added to counteract the effect of FGF19, which caudalises cells^7^. Initial comparison between conditions was performed on the H7 hESC line. qPCR was carried out on day 20, when cells are mostly neural precursors and day 50, when there are developing neurons. We did not find strong evidence that any of the 12 combinations of XAV939 and SHH significantly impacted the production of neural precursors on day 20 or neurons on day 50, although specific combinations had some effect on NESTIN and NEUN expression (Figure 1B and 1C): 200ng/ml of SHH decreased NESTIN expression in comparison to 0 ng/ml of SHH in the XAV939 2µM condition and 1µM of XAV939 decreased NEUN expression relative to 0µM of XAV939 when treated with 200ng/ml of SHH but not in any other conditions. Two specific combinations produced 2-6 times higher NKX2.1 expression than the rest: SHH 0 and 200ng/ml, both with XAV 1µM treatment (Figure 1D). This was not reflected in LHX6 expression on day 50, possibly meaning that some of the NKX2.1-expressing cells may have become LHX6-negative neurons, or cells are slowly progressing in their expression of identity, not yet expressing LHX6 in their full expression range. Within the fixed XAV concentration, there was no difference in LHX6 expression except for the highest XAV and SHH combination. When varying XAV939 with a fixed SHH concentration, the addition of XAV939 increased LHX6 expression in SHH 100 and 200ng/ml conditions (Figure 1E). As for PVALB and SST expression, the effect of different combinations of SHH and XAV939 was more striking. For PVALB, in the XAV 1µM condition, there was an increasing level of PVALB expression with increasing amount of SHH (Figure 1F), while for SST, lower SHH signalling produced higher amounts of SST in XAV 1 and 0µM conditions (Figure 1G). Based on these, we selected 4 conditions to investigate further: SHH 0 and 200ng/ml combined with 1 and 2µM XAV. These were named conditions 1 to 4: conditions 1 & 2 from XAV 1µM and conditions 3 & 4 from XAV 2µM (odd and even numbered conditions corresponding to SHH 0 and 200ng/ml, respectively).

**Figure 1.**
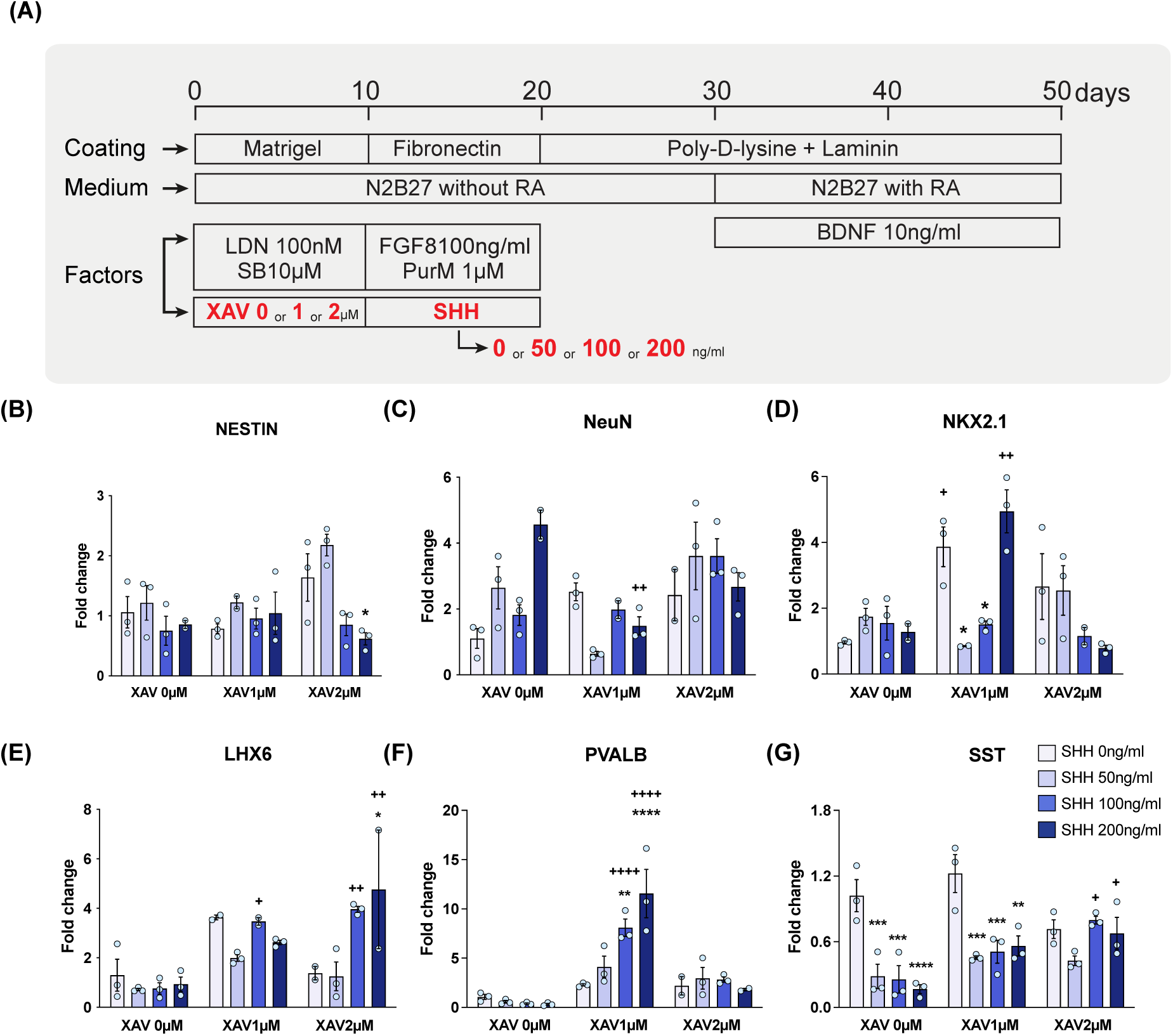
Comparison of 12 culture conditions to find the optimum condition to generate PVALB and SST cortical interneurons. (A)Experimental schematics. Day 0 is the human embryonic stem cell stage. (B)SHH significantly affected the expression of NESTIN (F_3,22_=5.243; P=0.0070), while XAV (F_2,22_=2.624; P=0.0950) and the interaction (F_6,22_=2.344; P=0.0667) did not. (C)XAV (F_2,21_=7.486; P=0.0035) and interaction (F_6,21_=5.137; P=0.0022) affected the expression of NEUN while SHH did not (F_3,21_=1.483; P=0.2481). (D)XAV (F_2,21_=7.168; P=0.0042) and interaction (F_6,21_=6.811; P=0.0004) affected the expression of NKX2.1 while SHH did not (F_3,21_=2.745; P=0.0686). (E)SHH (F_3,20_=4.533; P=0.0140), XAV (F_2,20_=16.41; P<0.0001) and interaction (F_6,20_=3.399; P=0.0178) affected the LHX6 expression. (F)SHH (F_3,22_=4.435; P=0.0139), XAV (F_2,22_=41.51; P<0.0001) and interaction (F_6,22_=7.118; P=0.0003) affected the PVALB expression. (G) SHH (F_3,24_=19.60; P<0.0001), XAV (F_2,24_=6.939; P=0.0042) and interaction (F_6,24_=4.257; P=0.0047) affected the SST expression. Two-way ANOVA with Bonferroni-adjusted multiple comparison test was performed. Each dot is a biological repeat with a minimum of two per condition. *, p<0.05, **, p<0.01, ***, p<0.001, ****, p<0.0001 compared to SHH 0ng/ml within the same XAV treatment. +, p<0.05, ++, p<0.01, ++++, p<0.0001 compared to XAV 0µM within the same SHH treatment.

### Single-cell experiments confirm the one condition that favours PVALB expression

The chosen conditions were further tested in H7 plus two additional hPSC lines (H9 hESCs and IBJ4 iPSCs) to verify the efficacy of the protocol. Firstly, cells were harvested on day 50 for bulk qRT-PCR. As with our previous experiment, all three cell lines were consistent in producing the highest amount of PVALB under condition 2 (XAV 1 µM with SHH 200 ng/ml), with the H9 hESC line producing the highest level of PVALB expression (Figure 2C). LHX6 expression showed a similar level across all cell lines, apart from condition 4 in IBJ4 iPSC lines (Figure 2A). SST expression patterns differed between cell lines: H7 showed the highest SST expression in condition 1; H9 in condition 2; and IBJ4 in condition 4. Overall, condition 2 appeared to produce the highest amount of PVALB, together with either the highest or average levels of SST. We then performed fluorescence *in situ* hybridisation (FISH) using RNAscope^TM^ probe for LHX6, SST and PVALB to quantify the percentage of cells expressing these markers in H9. Neither LHX6 nor SST showed a significant difference between the four conditions (Figure 2D, 2E, 2H and 2I). However, PVALB expression was significantly higher in condition 2, which is in line with the qRT-PCR result (Figure 2F, 2G and 2J). The percentage of PVALB-expressing cells was approximately 10% in condition 2, roughly double that seen in other conditions. We further validated the protocol via single-cell qRT-PCR using the Fluidigm C1 platform. In line with the FISH result, we found PVALB expression was highest in condition 2, with around 10% of the total population being positive (Figure 2K). Finally, we investigated the protein expression of PVALB and SST. For PVALB, condition 2 again showed the highest expression, although the absolute percentage was not as high as mRNA expression (2% vs 10%, as protein expression often lags behind mRNA expression^11^) (Figure 2L and 2R). Mirroring the bulk qRT-PCR data, SST protein was most abundantly expressed in condition 2 (Figure 2M and 2R). We also found that the majority of cells in condition 2 expressed ventral forebrain markers such as FOXG1 and NKX2.1 (Figure 2N and 2O) and progenitor marker SOX2 on day 20 (Figure 2P) and GABA on day 50 (Figure 2Q). These experiments demonstrate that, of the four conditions tested, condition 2 is the best for producing PVALB-expressing cortical interneurons.

**Figure 2.**
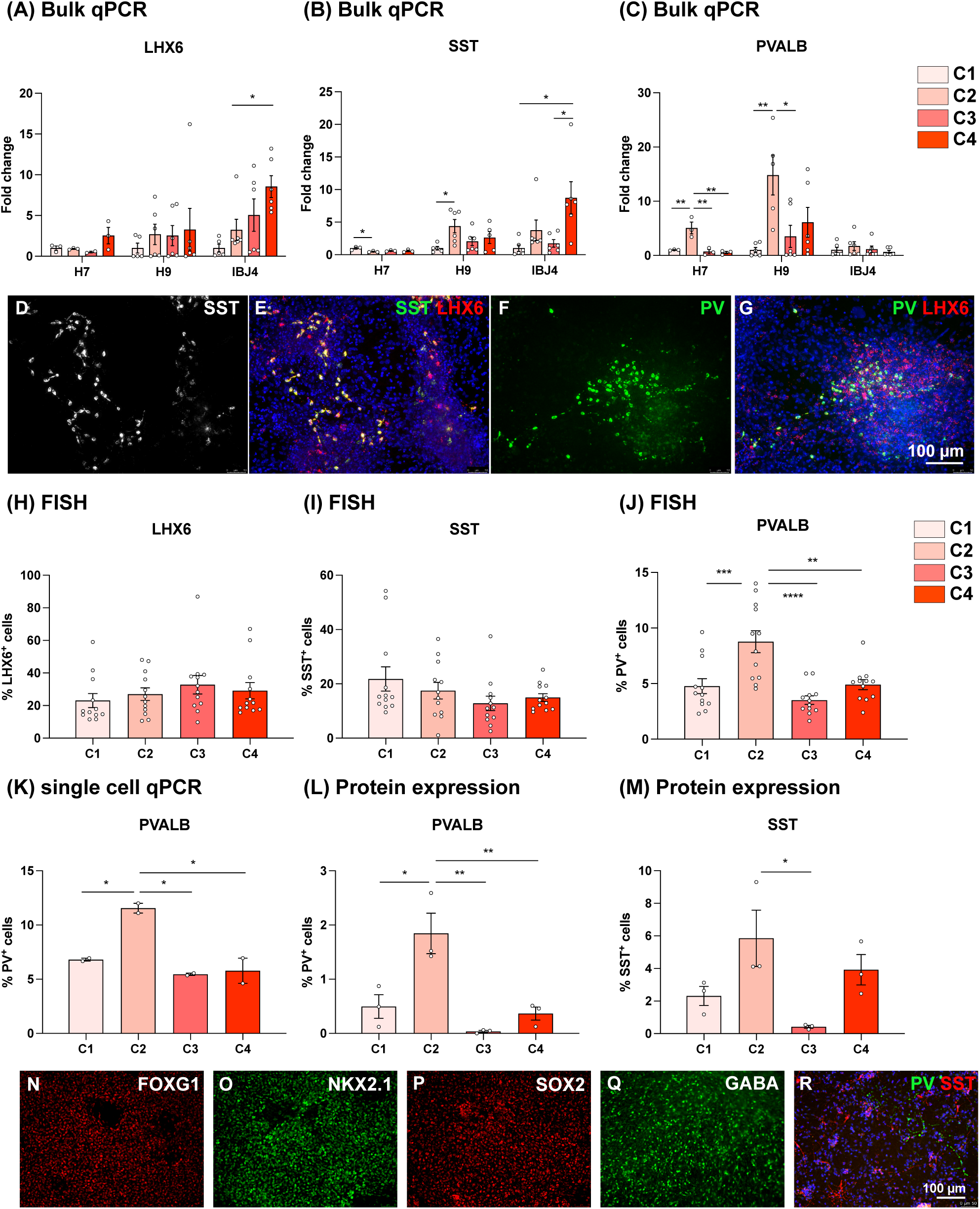
Molecular analysis of *in vitro*-generated cortical interneurons. Bulk qPCR analysis of LHX6, SST and PVALB on three hPSC lines on differentiation day 50. (A)LHX6 expression was significantly different in IBJ4 (F_3,19_=4.708; P=0.0127) across 4 conditions, but was not in H7 (F_3,7_=2.350; P=0.1587) and H9 (F_3,19_=0.3100; P=0.8179). (B)SST expression was significantly different across 4 conditions in H7 (F_3,8_=4.837; P=0.0332), H9 (F_3,18_=3.222; P=0.0473) and IBJ4 (F_3,20_=5.231; P=0.0079). (C)PVALB expression was significantly different in H7 (F_3,8_=13.76; P=0.0016) and H9 (F_3,19_=5.815; P=0.0054) but not in IBJ4 (F_3,20_=0.6193; P=0.6107). (D-G)FISH using SST, PVALB and LHX6 probes on day 50. (H-J)LHX6 (F_3,44_=0.7324; P=0.5382) and SST (F_3,44_=1.543; P=0.2168) expression were not different across the 4 conditions, but PVALB expression was (F_3,44_=11.62; P<0.0001). (K)PVALB expression was significantly higher in condition 2 (F_3,4_=20.24; P=0.0070). (L)PVALB (F_3,8_=12.58; P=0.0021) and (M)SST (F_3,8_=5.085; P=0.0293) expression was significantly higher in condition 2. One-way ANOVA with Bonferroni-adjusted multiple comparison test was performed. Each dot is a biological repeat with a minimum of two per condition. *, p<0.05, **, p<0.01, ***, p<0.001, ****, p<0.0001 compared to C1. D-M and R are H9 and N-Q are H7.

### Single-cell RNA sequencing analysis confirms the generation of authentic PVALB-expressing cortical interneurons from hPSCs

While previous 2D differentiation studies have reported evidence suggestive of PVALB interneuron generation, none have successfully identified PVALB-expressing interneuron populations in single-cell RNA sequencing (scRNAseq) data^5–7,12^. We, therefore, performed a scRNAseq experiment to fully analyse the differentiated cells from hPSCs using our chosen condition 2. H7 hESCs were differentiated, harvested at days 20, 30 and 50, and processed with 10x genomics Chromium Single Cell 3ʹ Reagent Kits v3. We aimed to capture 3,000 cells per sample and sequence 50,000 reads/cell using Illumina HiSEq4000. Cells were filtered based on % of mitochondria genes (<20%) and number of genes & counts (>5% and <95%) to filter out dead cells and doublets (Figure S2A and S2B). We obtained 9044 cells with 20,688 features/genes. Cells were clustered in Monocle3, and UMAP was used for visualisation. This initial analysis yielded 27 clusters (Figure S2C and S2D). Clusters were further evaluated and quality-checked using gene counts to drop low-quality clusters (Figure S2E), resulting in a final set of 21 clusters (Figure 3A and 3B) for 7961 cells, each containing, on average, 4158 genes per cell. As cells became more developed/differentiated, they formed more diverse populations, as demonstrated by a larger number of clusters containing day 50 cells than containing day 30 or day 20 cells. To determine cluster cell type identity, we first identified a set of marker genes for each cluster: genes that were differentially expressed when comparing cells within the cluster to cells outside the cluster. Marker sets were identified in a similar fashion for known inhibitory/progenitor cell types from human foetal (5.85 to 37 post-conception week) cortex and ganglionic eminence using published scRNAseq data^13^. To determine which cell type each *in vitro* cluster most resembled, we tested cluster marker sets for enrichment with *in vivo* cell type markers using a Fisher’s Exact test (Figure 3C and Figure S2F). Clusters highly enriched for *in vivo* markers of newborn and early developing inhibitory neurons (NewN, DevN) were co-localised in the UMAP plot, clearly separated from those enriched for *in vivo* progenitor markers, with clusters possessing dual progenitor-neuron enrichment (TransRG) – indicating cells transitioning from the former to the latter – located in between (Figure 3A, 3B and 3C). A branch of clusters corresponding to dividing intermediate progenitors (IPC) could also be identified, emerging from the top right-hand corner of the main body of progenitors. The remaining clusters comprised radial glia (RG), which were organised into 4 groups based on *in vivo* marker enrichment (Figure 3C). These cluster identities were in line with the canonical marker gene expression. Clusters showed telencephalic identity (FOXG1, SIX3, HES5) with majority being neural precursor cells (SOX2, NES) while dorsal telencephalic (PAX6, DMRT5, EMX2, TBR1), diencephalic (DBX1, FOXB1, GBX2, RAX), mid and hindbrain (EN2, GBX2, IRX3, PAX2) genes were minimally seen (Figure 3D and Figure S3), demonstrating ventral telencephalic fate of the cells. Neuronal clusters expressed canonical neuronal genes such as DCX, MAP2, STMN2 and MAPT, while more mature neuronal gene NEUN (RBFOX3) was expressed only in maturer cells (Figure 3E). Synaptic genes such as SYP and DLG4, and GABAergic markers such as GAD1 and GAD2 were also expressed in neuronal clusters together with migrating interneuron marker ARX (Figure 3F), demonstrating inhibitory neuronal lineage. MGE and cortical interneuron genes were also expressed. The majority of cells expressed NKX2.1 and MAF (MGE progenitor genes), and neuronal clusters expressed the cortical interneuron transcription factor SOX6 and LHX6 (although less amounts) (Figure 3G). SST and PVALB mRNA were also expressed in neuronal clusters, with strong SST expression seen in the more developed side of clusters (DevN clusters 9, 10 and 12, with higher expression further away from NewN). As expected, fewer PVALB-expressing cells were seen, and these were largely restricted to the less developed neuronal clusters (NewN cluster 14 and 21). This is in line with their temporal difference in birthdate: SST neurons are born earlier than PVALB neurons^14^; hence, at the same timepoint, SST neurons will be in a more developed state.

**Figure 3.**
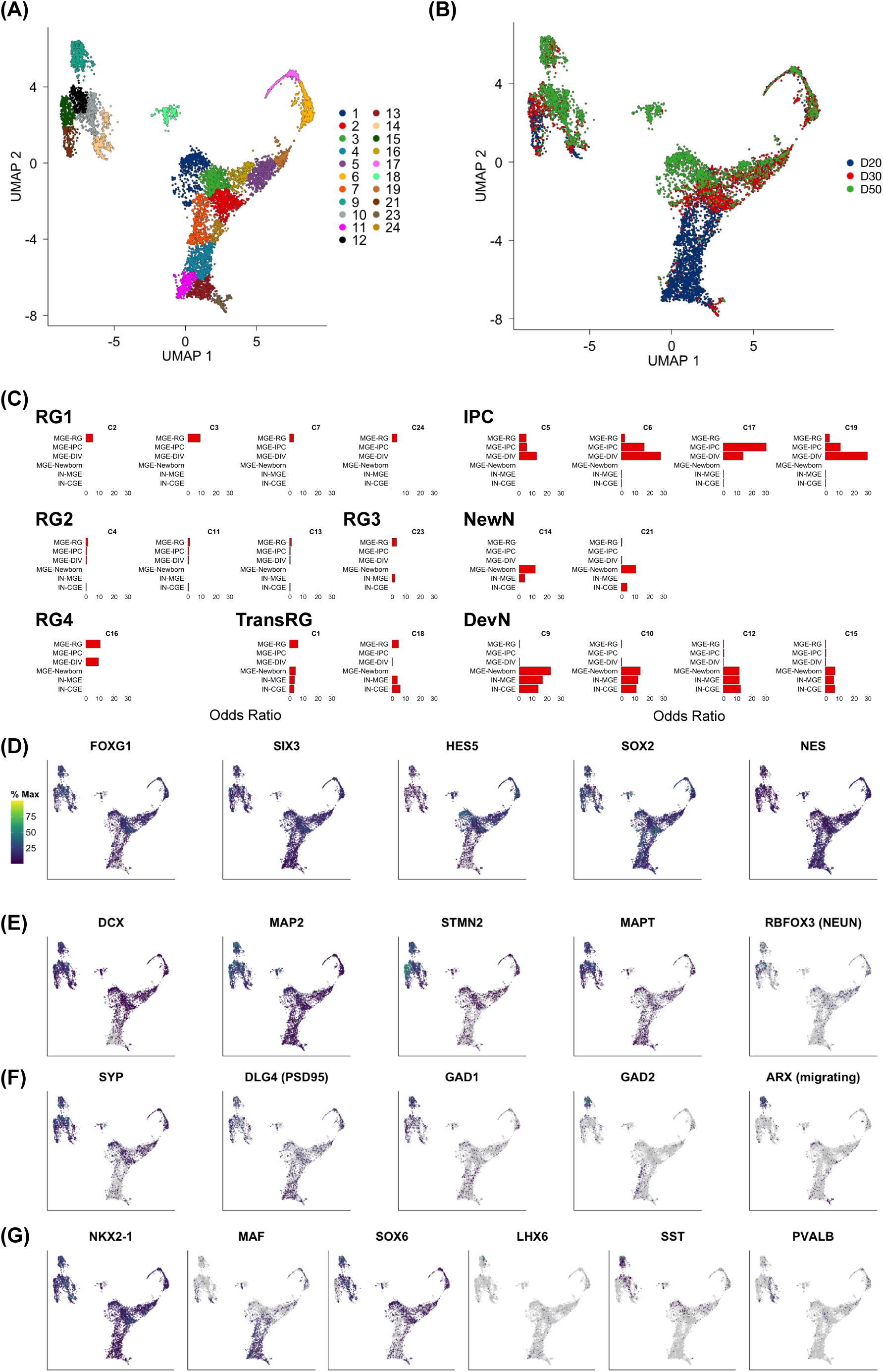
Single cell transcriptome analysis on day 20, 30 and 50 cultures differentiated using condition 2. (A)UMAP showing 21 clusters (B)Culture age of each cluster (C)Enrichment of DEGs from known *in vivo* clusters. Expression of telencephalic (D), neuronal (E), synaptic and GABAergic (F) and MGE (G) genes.

To explore the identity of neurons generated using our condition 2 protocol in more detail, we compared our data to a scRNAseq dataset from foetal, postnatal and adult human cortical tissue, within which interneuron sub-types had been identified^15^ (Figure 4A and 4B). We first combined our neuronal clusters (9, 10, 12, 14, 15 and 21) with the interneuron clusters identified by Velmeshev et al.^15^ and reclustered using Seurat-CCA. Our cells clustered together with those of Velmeshev et al. (Figure 4C). Most cells were located in the centre of the plot, although two groups of cells clearly overlapped *in vivo* SST and PVALB clusters.

**Figure 4.**
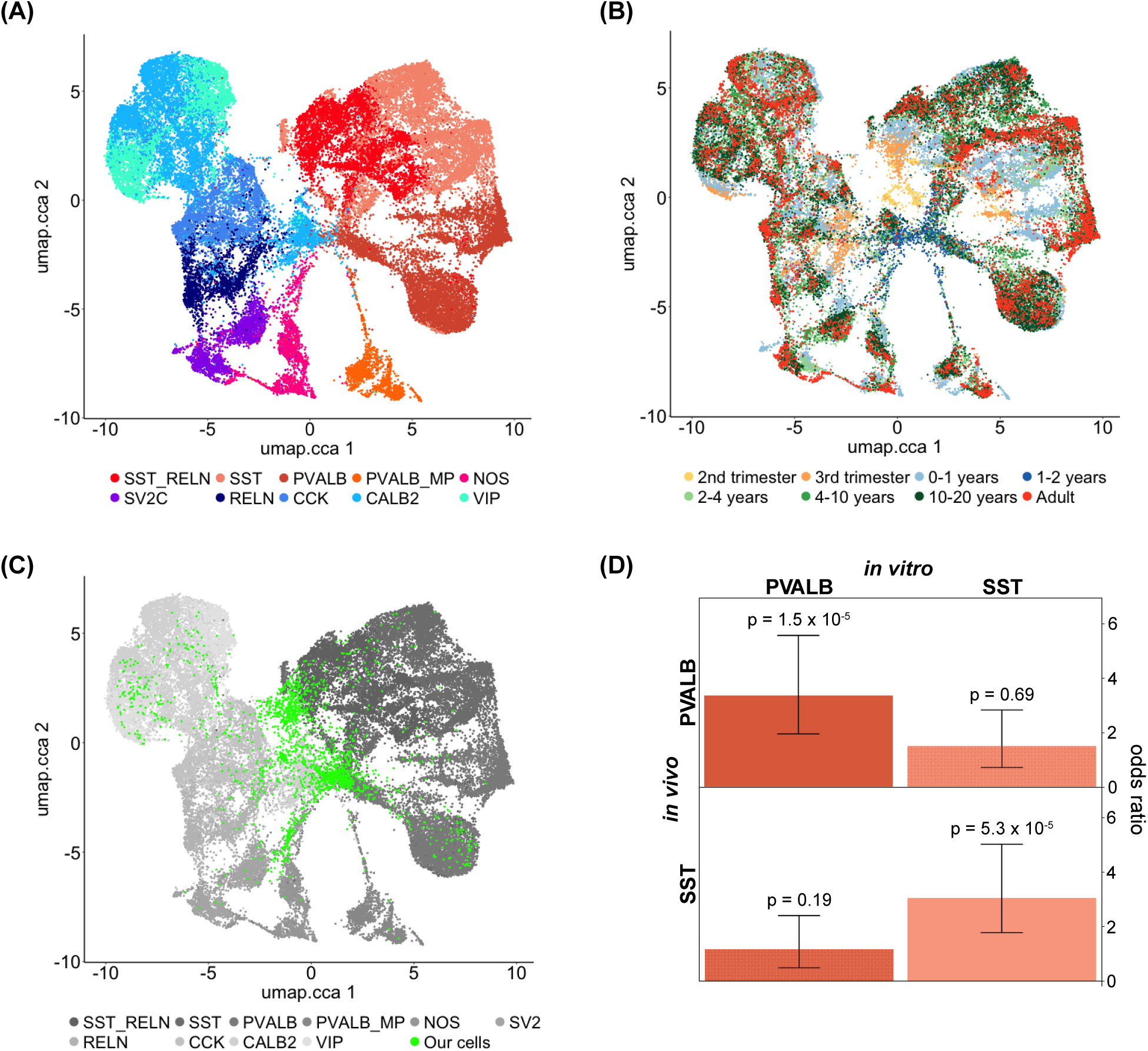
Comparison to the single-cell transcriptomes of *in vivo* cortical interneurons. (A) *in vivo* data set. (B)Sample ages (C)Co-clustering with *in vitro*-generated and *in vivo* cortical interneuron single-cell transcriptomes. (D)Enrichment of *in vivo* PVALB and SST neuronal genes in *in vitro*-generated PVALB and SST cortical interneurons.

Finally, we directly compared gene expression patterns between our most developed SST and PVALB neurons and their *in vivo* counterparts. Taking the cells from clusters with the highest, most concentrated PVALB (C10, C14) and SST (C9, C12) expression (Figure 3), we performed differential expression analysis to identify genes up-regulated in PVALB relative to SST cells *in vitro* (and vice versa). Using the Velmeshev et al.^15^ data, genes up-regulated in PVALB and SST cells *in vivo* were identified in the same fashion. Using Fisher’s exact test to evaluate the similarity between *in vitro* and *in vivo* expression (Figure 4D), PVALB up-regulated genes identified *in vitro* were highly enriched for PVALB (P_Bonferroni_=6.07x10^-5^) but not SST (P_Bonferroni_=0.74) up-regulated genes identified *in vivo*. Similarly, SST up-regulated genes identified *in vitro* were highly enriched for SST (P_Bonferroni_=2.12x10^-4^) but not PVALB (P_Bonferroni_=1) up-regulated genes identified *in vivo*. These data demonstrate that our optimised 2D interneuron protocol generates authentic cortical PVALB and SST interneurons that have similar transcriptional signatures to those in the human brain.

## Discussion

Our comparison of 12 culture conditions, testing different combinations of SHH and a WNT inhibitor (XAV939), identified a condition that produces a substantial number of PVALB- and SST-expressing cortical interneurons *in vitro* within 50 days of directed differentiation from human pluripotent stem cells. In contrast to all previously published 2D *in vitro* protocols, we were able to identify PVALB-expressing cortical interneurons in single-cell transcriptomic data. The study by Close *et al*^12^ sequenced transcriptomes of 1700 single cells over 4 different culture days but found no PVALB-expressing cells. This was interesting as they have followed the protocol of one of the earlier studies, which showed the presence of PVALB protein-expressing neurons^5^. The discrepancy could be due to the non-specificity of the antibody used to detect PVALB cells and the varying maturation status of the neurons resulting from the use/absence of mouse cortical feeders. In addition, this was not the sole case that showed discrepancies. Kim *et al*^7^ showed the expression of PVALB in their *in vitro* differentiated interneurons in a 2014 study (0.87% of *in vitro* cells by week 6); however, their following studies focusing on identifying disease mechanisms^16,17^ did not detect

PVALB-expressing interneurons *in vitro* in either transcriptomic or staining analyses. The discrepancy could be due to the low level of PVALB neuron generation in the original protocol and variation in the differentiation response from different hPSC lines to the protocol, as the latter studies used different lines to the initial study (patient/healthy volunteer-derived iPSCs). In contrast, we now provide a protocol that generates a substantial population of PVALB-expressing cortical interneurons (∼10% of day 50 cells) and performs consistently across multiple hPSC lines.

SHH and WNT play a key role in determining the regional identity of the developing brain. High WNT expression is seen in the caudal and dorsal part of the developing brain^18^, promoting pallial identity^19^, while high SHH expression is seen in the ventral part^20^, specifying MGE identity^21^. Accordingly, promoting SHH and counteracting WNT signalling has been the strategy for generating cortical interneurons. However, the careful optimisation of these two molecules has not been conducted so far, creating a non-ideal MGE environment to generate diverse cortical interneuron subtypes, especially the PVALB-expressing subtype. Previous studies utilised a range of different small molecules to target the SHH and WNT pathways, making direct comparison with our conditions challenging. However, the studies utilising XAV939 (WNT inhibitor) employed higher concentrations in general (2 µM)^5,12,22^ compared to ours (1 µM). We tested 2 µM in our 12 conditions, but it resulted in lower PVALB expression. Interestingly, Bershteyn *et al*^23^ used a much higher concentration, 10 µM of XAV939, and their cells *in vitro* did not produce PVALB mRNA. Another factor to consider is the use of mitotic inhibitors. Studies aiming to produce a pure MGE/cortical interneuron population for transplantation purposes utilised mitotic inhibitors^23,24^ such as DAPT (NOTCH inhibitor), PD0325901 (MEK inhibitor) and PD0332991 (CDK inhibitor), including when analysing cells *in vitro*. These inhibitors induce synchronised cell cycle exit and may be suitable for preparing homogenous neuronal donor cells. However, the PVALB-expressing interneurons are predominantly generated from basal progenitors (intermediate precursor cells) in the subventricular zone, which are derived from apical progenitors (radial glia), while SST-expressing interneurons are directly generated from apical progenitors in the ventricular zone^25^. The use of inhibitors may have impeded the generation of basal progenitors, thereby reducing the generation of PVALB-expressing interneurons. Within MGE, SST and PVALB-expressing neurons are preferentially generated from dorsal and ventral parts, respectively^9,26^ with higher SHH signalling reported in the dorsal compared to the ventral MGE^10^. Under our XAV 1µM condition, increasing the concentration of exogenous SHH resulted in elevated PVALB mRNA levels in H7 (Figure 1F), H9 and IBJ4 (Figure 2C). In contrast, SST expression showed a less consistent pattern. In H7, lower SHH treatment led to higher SST mRNA levels (Figure 1G), whereas in H9 and IBJ4, higher SHH concentrations increased SST expression under either XAV 1µM or 2µM conditions (Figure 2B). Notably, the observed PVALB expression pattern is the opposite of what would be expected based on *in vivo* dorsal-ventral MGE patterning. One possible explanation is that the timing of SHH exposure in our protocol corresponds to an earlier or broader dorsal-ventral patterning window. In this context, higher SHH levels may promote ventral forebrain identity rather than dorsal MEG fate. Consistent with this interpretation, the previous study showing preferential generation of SST over PVALB-expressing neurons in ventral MGE applied exogenous SHH after cells had already acquired a ventral telencephalic identity^10^. FISH analysis suggests an additional possibility. Although our optimised condition 2 yielded the highest proportion (10%) of PVALB-expressing cells, this remained lower than the proportion of SST-expressing cells (∼20%) (Figure 2I and 2J). A similar trend was observed at the protein level, with approximately 2% PVALB-positive cells compared to 6% SST-positive cells. Together, these findings suggest that our optimised condition may preferentially generate dorsal MGE rather than ventral MGE identity, producing both interneuron subtypes but with a bias toward SST. Further refinement of SHH treatment timing and dosage may better recapitulate ventral MGE-like conditions and increase the relative production of PVALB-expressing interneurons.

PVALB-expressing interneurons have a distinct electrophysiological property – high frequency firing (∼200Hz)^27^. While our cells have the ability to fire *in vitro*, they do not display high-frequency firing (data not shown); this may be due to their immature status. Other studies^23,24,28^ demonstrated functional fast spiking PVALB-expressing interneurons 14 months after transplantation into rodent brains, implying that a long time period is required for these interneurons to become mature. It is also known that these cortical interneurons mature postnatally in the rodent brain^29,30^ and presumably in human brains, too. Therefore, it is not surprising to observe the lack of high-frequency firing in our cells. To fully capture the electrophysiological properties of our *in vitro*-generated cortical interneurons, transplantation will be required.

The origin and timing of cortical interneuron fate decision is thought to be either at the late progenitor stage in MGE or after tangential migration into the cortex, with both hypotheses finding support in the literature^31,32^. Our results suggest that having the right type of progenitors is important for producing PVALB-expressing interneurons, as our culture conditions are different until cells reach the progenitor stage from the pluripotent stage (i.e. the same culture medium is used once cells reach the progenitor stage). This indicates that the fate of the PVALB-expressing interneurons in our culture is already specified at the progenitor stage. Having said that, it will be important to analyse our progenitor populations further to identify possible PVALB-fated progenitor groups. A mouse study^33^ found that MAF is highly expressed in progenitors that will later become SST-expressing interneurons. Some of our RG clusters (C4, C11, C13 and C23) express MAF at especially high levels, and further lineage tracing experiments would allow us to reveal whether these groups of progenitors produce SST-expressing interneurons, and whether groups negative for MAF generate PVALB-expressing interneurons.

Our optimised cortical interneuron differentiation protocol enables the generation of PVALB- and SST-expressing cortical interneurons within 50 days of *in vitro* differentiation from the pluripotent stage, without the need for forced gene expression, cell sorting, mitotic inhibitors, or rodent primary co-culture. A recent assembloid study^34^ combining dorsal and ventral forebrain organoids reported the emergence of functional PVALB interneurons after 120 days of 3D culture. Although such approaches are valuable, 3D cultures exhibit increased batch-to-batch variability and require more time- and resource-intensive analyses. While the proportion of PVALB-expressing neurons generated here is moderate (∼10% mRNA expression), consistent with the prolonged maturation timeline of cortical interneurons, this protocol represents a meaningful advance. It enables the study of human PVALB interneuron development entirely *in vitro* using a 2D system, circumventing the variability associated with animal models and 3D cultures, and providing a robust foundation for adoption and further optimisation. Future studies extending this platform may advance our understanding of PVALB interneuron fate specification, functional maturation, and disease-relevant mechanisms.

## Methods

### Key Resources Table

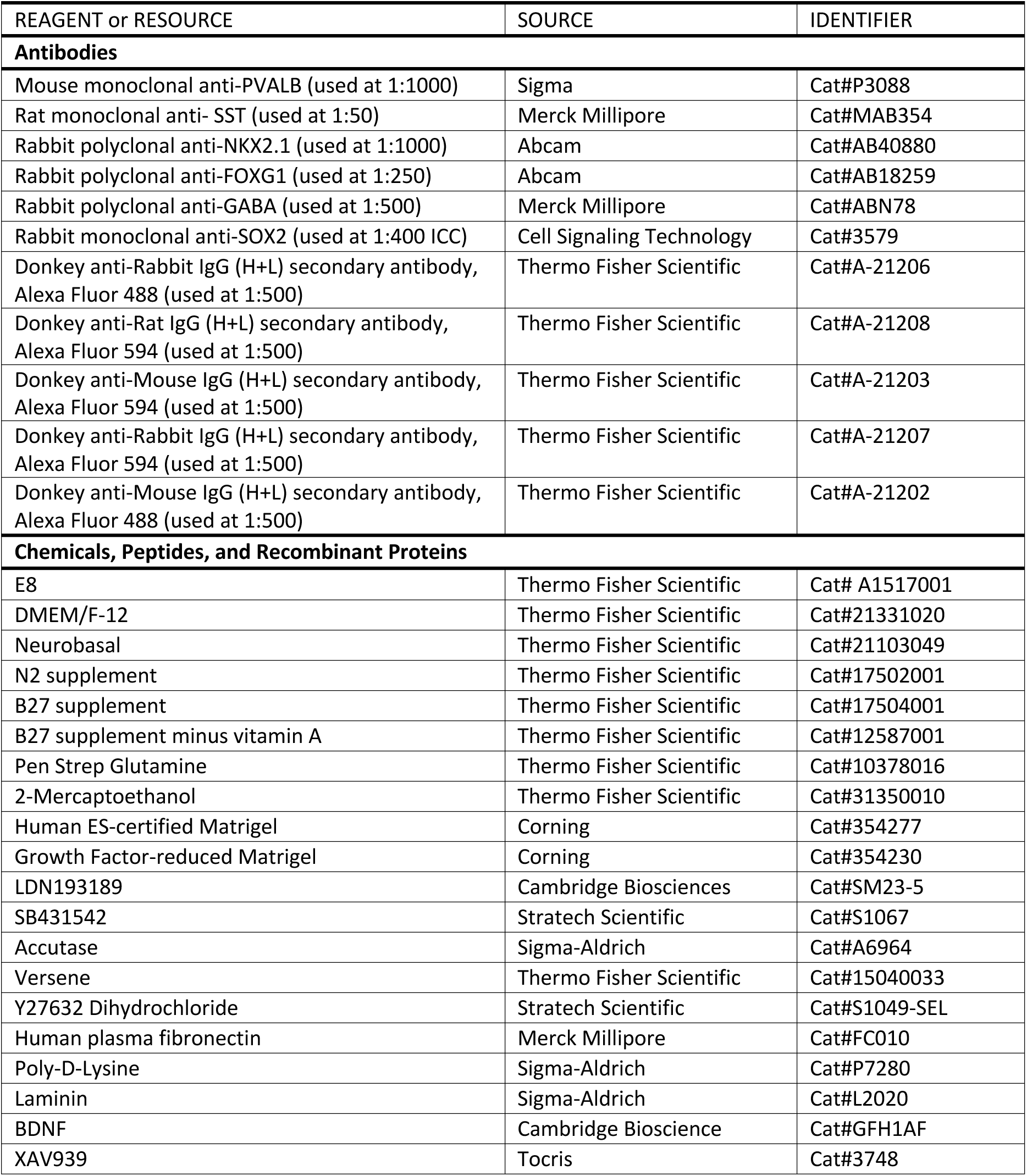
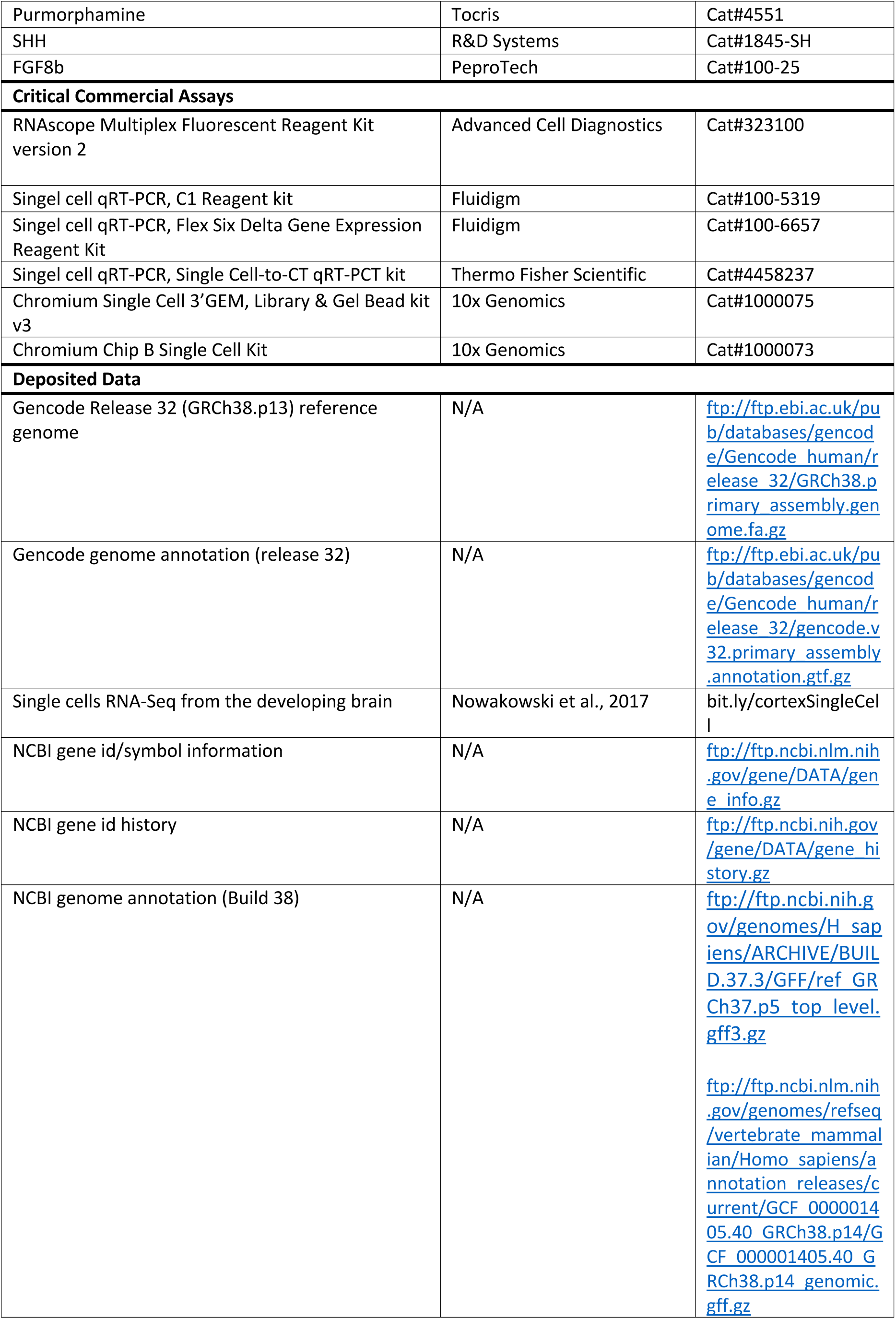
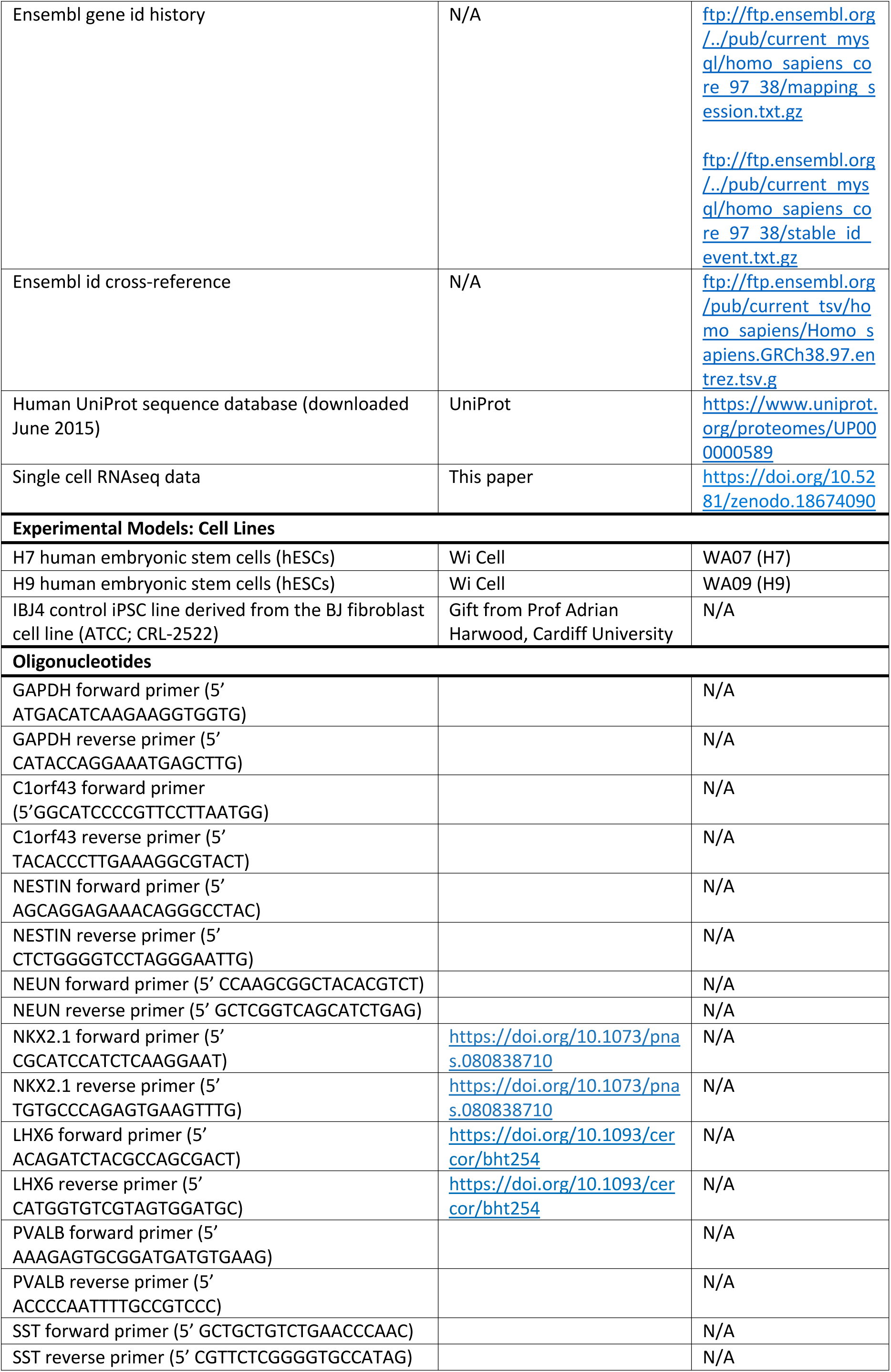
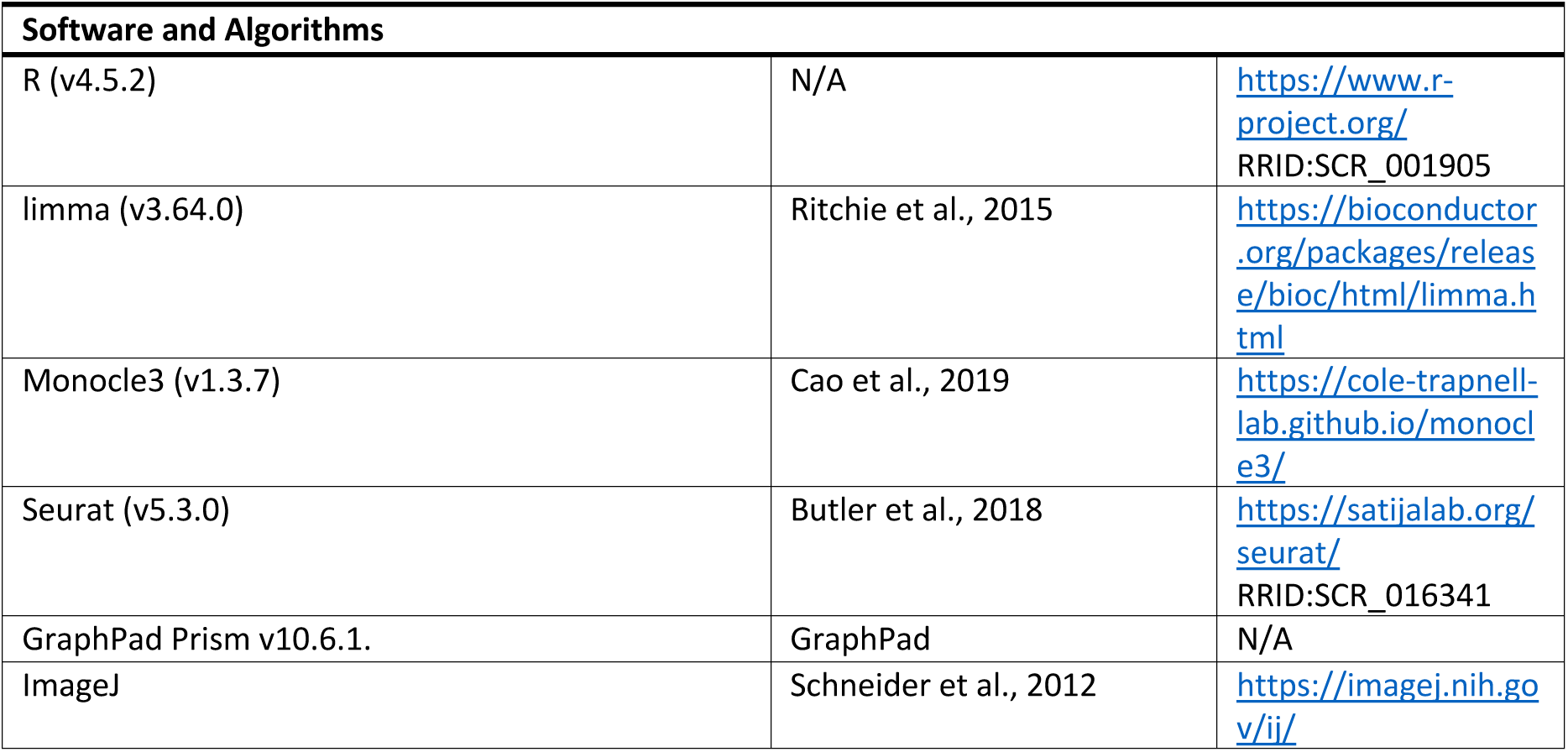

### hPSC culture

All hPSC lines were maintained at 37°C and 5% CO_2_ in 6-well cell culture plates (Greiner) coated with 1% Matrigel hESC-Qualified Matrix (Corning) prepared in Dulbecco’s Modified Eagle Medium: Nutrient Mixture F-12 (DMEM/F12, Thermo Fisher Scientific). Cells were fed daily with mTeSR1 (STEMCELL Technologies) and passaged at 80% confluency using Versene solution (Thermo Fisher Scientific) for 1.5 minutes at 37°C followed by manual dissociation with a serological pipette.

### Cortical interneuron differentiation

Differentiation to cortical interneurons (Figure 1A) was achieved using the dual SMAD inhibition protocol^35^ with the addition of WNT inhibitor (XAV939), SHH and its agonist (Purmorphamine) and FGF8. Prior to differentiation, Versene treatment and mechanical dissociation was used to passage hPSCs at approximately 100,000 cells per well into 12-well cell culture plates coated with 1% Matrigel Growth Factor Reduced (GFR) Basement Membrane matrix in DMEM/F12. Cells were maintained in mTeSR1 at 37°C and 5% CO_2_ until 90% confluent. At day 0 of the differentiation, mTeSR1 media was replaced with N2B27-RA neuronal differentiation media consisting of: 2/3 DMEM/F12, 1/3 Neurobasal, 1x N-2 Supplement, 1x B27 Supplement minus vitamin A, 1x Pen Step Glutamine and 50 µM 2-Mercaptoethanol, which was supplemented with 100nM LDN193189, 10µM SB431542 and XAV939 (0, 1 and 2µM) for the first 10 days only (the neural induction period). At day 10, cells were passaged at a 2:3 ratio into 12 well cell culture plates coated with 15µg/ml Human Plasma Fibronectin in Dulbecco’s phosphate-buffered saline (DPBS). Passage was as previously described, with the addition of a 1-hour incubation with 10µM Y27632 Dihydrochloride (ROCK inhibitor) prior to Versene dissociation. During days 10 to 20 of differentiation, cells were maintained in N2B27-RA with FGF8 100ng/ml, Purmorphamine 1µM and SHH 0, 50, 100 or 200ng/ml and passaged at day 20 in a 1:2 ratio into 24-well cell culture plates, sequentially coated with 10µg/ml poly-d-lysine hydrobromide (PDL) and 15µg/ml laminin in DPBS. Together with 10ng/ml of BDNF, Vitamin A was added to the differentiation media at day 30, with standard 1x B27 Supplement (Thermo Fisher Scientific), replacing 1x B27 Supplement minus vitamin A, and cells were maintained in the resulting N2B27+RA media for the remainder of the differentiation. For the entire differentiation period, half of the medium was replenished every other day.

### Bulk qRT-PCR

At days 20 and 50, cells were washed with DPBS and lysed with 1mL TRIzol per well. Lysates were mixed with 200μl chloroform and centrifuged at 12,000g for 15min at 4°C. RNA in the aqueous phase was precipitated with 0.5ml of isopropanol, washed and dried with 75% ethanol, and resuspended in 30μl RNase-free water. Genomic DNA was removed with PerfeCTa DNase (Quanta), and cDNA was synthesised with qScript cDNA Supermix (Quanta). qPCR was performed using iTaq™ Universal SYBR® Green Supermix (BioRad) in technical and biological duplicates/triplicates in a CFX Connect System (BioRad) under the following conditions: 94°C for 4min; 39 cycles of 94°C for 30s, 60°C for 15s, 72°C for 30s. Gene expression was calculated using the 2^(–ΔΔCt) method, normalised to the average of GAPDH and C1orf43, and calibrated to a specified control condition.

### Fluorescent *in situ* hybridisation

NPCs (day 18-19) derived from H9 were cultured at a density of 20,000 cells per well of a 96-well plate. At day 50, cells were fixed in 4% paraformaldehyde (PFA) for 30min. The RNAscope® Multiplex Fluorescent Assay v2 was performed according to the manufacturer’s instructions for adherent cells. Samples were hybridised with target probes against *SST* (Probe-C1), *LHX6* (Probe-C2), and *PV* (Probe-C3). Probes were detected using HRP-based amplification and conjugated fluorophores (Cy3 for *SST*, Cy5 for *LHX6*, and Fluorescein for *PV*). Nuclei were counterstained with DAPI. Z-stack images were acquired from five random fields per condition using a 20x objective on an inverted Leica DMI6000B microscope. Positive cells for each marker were quantified manually, while DAPI+ nuclei were counted using Cell Profiler software.

### Single-cell qRT-PCR

H9 hESCs were differentiated under four conditions (C1–C4) with two biological replicates per condition. Single-cell gene expression was assessed at neuronal maturity (day 50 onwards); due to scheduling constraints, samples were processed between days 57–70 with even sample distribution on each day. Cultures were treated with 10μM Y-27632 for 1–2hr prior to dissociation. Cells were washed with PBS and dissociated using Accutase (5-10min, 37°C), collected in N2B27 supplemented with RA and BDNF, centrifuged (200 × g, 5min), filtered, and counted. Cells were mixed with Fluidigm suspension reagent (3:1 ratio) and adjusted such that 6μl contained ∼1000 cells. Single cells were captured and pre-amplified using a C1 Single-Cell PreAmp IFC (10–17μm; Fluidigm). Wells containing multiple or damaged cells were excluded following microscopic inspection. Capture efficiency ranged from 68–97%. Gene expression (GAPDH, C1orf43, PV) was quantified using a BioMark HD system with a Flex Six IFC chip (GE Flex Six PCR + Melt v1). Amplification and melting curves were manually inspected, and aberrant profiles were excluded. After quality control, 116 (C1), 147 (C2), 118 (C3), and 140 (C4) single cells were analysed.

### Immunocytochemistry

Cells were fixed in 4% PFA in PBS for 20min at 4°C, followed by a 1-hour room temperature incubation in blocking solution of 5% donkey serum (Biosera) in 0.3% Triton-X-100 (Sigma) PBS (0.3% PBST). Primary antibodies, used at an assay-dependent concentration, were diluted in blocking solution and incubated with cells overnight at 4°C. Following the removal of the primary antibody solution and three PBS washes, cells were incubated in the dark for 2hr at room temperature with the appropriate Alexa Fluor secondary antibodies, diluted 1:500 in blocking solution. After an additional two PBS washes, cells were counterstained with DAPI nucleic acid stain, diluted 1:1000 with PBS, for 5min at room temperature and following a final PBS wash, mounted using Dako Fluorescence Mounting Medium (Agilent) and glass coverslips. Imaging was with either the Zeiss LSM710 confocal microscope or the Leica DMI6000B inverted fluorescent microscope. In each well, images were taken from four to five random fields. Images were Z-stacked and processed on the LAS-X software. The cells were counted manually with ImageJ to detect the expression of SST and PVALB.

### Single Cell 3ʹ gene expression libraries preparation

At days 20, 30 and 50 of differentiation, 4800 single cells and 42.6μl of Nuclease-free Water were prepared to add to the 33.4μl of Master Mix, following Step 1.1 Prepare Master Mix p24 of Chromium Single Cell 3ʹ Reagent Kits v3 User Guide | Rev A. Subsequently, 75μl of this preparation was loaded into the Chromium Chip B following Step 1.2 Load Chromium Chip B p25 of Chromium Single Cell 3ʹ Reagent Kits v3 User Guide | Rev A. The rest of the method for generating Single Cell 3ʹ gene expression libraries was carried out in exact accordance with the manufacturer’s protocol (Chromium Single Cell 3ʹ Reagent Kits v3 User Guide | Rev A).

### Sequencing

The single-cell libraries containing 75-bp paired-end reads were then delivered to the sequencing facility at the MRC Centre for Neuropsychiatric Genetics and Genomics at Cardiff University, and sequenced on the Illumina® HiSeq4000 across five lanes on one flowcell. The sequencing was performed exactly according to the manufacturer’s instructions (see 3ʹ Gene Expression Library Sequencing Depth & Run Parameters, library loading and pooling in page 46 of Chromium Single Cell 3ʹ Reagent Kits v3 User Guide | Rev A).

### Single-cell/nucleus RNA sequencing data

H7 hESCs were differentiated, harvested at days 20, 30 and 50, and processed with 10x genomics Chromium Single Cell 3ʹ Reagent Kits v3. It was planned to achieve 3,000 cells per sample and sequence 50,000 reads/cell using Illumina HiSEq4000. First, only features detected in at least 10 cells and cells with at least 200 detected features were retained for analysis. Then, cells were filtered based on % of mitochondria genes (<20%) and number of genes & counts (>5% and <95%) to filter out dead cells and doublets. We obtained 20,688 features/genes across 9,044 samples/cells. Cells were clustered in Monocle3 using the standard PCA method with 100 principal components, and UMAP was used for visualisation. This initial analysis yielded 27 clusters. Clusters were further evaluated and quality-checked using the number of genes and counts to drop low-quality clusters, resulting in a final set of 21 clusters for 7,961 cells. Marker genes for each cluster were identified using the FindAllMarkers function in Seurat (v5.3.0), applying standard parameters and comparing cells within each cluster to all cells outside that cluster. Differentially expressed genes were considered significant if their p-value after Bonferroni correction was <0.05.

Single-cell RNASeq gene expression data from (Nowakowski et al., 2017)^13^ were downloaded from bit.ly/cortexSingleCell. Cells corresponding to distinct neurodevelopmental cell-types (including cell-types at different stages of maturity in PFC and V1 cortex and MGE) were identified and extracted, collating all cells from the corresponding *in vivo* cell clusters as follows:

IN_CGE: “IN-CTX-CGE1”, “IN-CTX-CGE2”;

IN_MGE: “IN-CTX-MGE1”, “IN-CTX-MGE2”;

MGE_Newborn: “nIN1”, “nIN2”,“nIN3”, “nIN4”, “nIN5”;

MGE_IPC: “MGE-IPC1”,“MGE-IPC2”,“MGE-IPC3”;

MGE_RG: “MGE-RG1”,“MGE-RG2”;

MGE_DIV: “MGE-div”.

The resulting dataset consisted of 974 cells and 17,764 protein-coding genes. The Limma package (v 3.64.0) was used to identify differentially expressed genes comparing cells within each cluster to all cells outside that cluster. Differentially expressed genes (DEGs) were considered significant if they had a fold change > 2 and their p-value after Bonferroni correction was <0.05.

Single-cell RNA-seq gene expression data and metadata from Velmeshev et al. 2023^13^ were downloaded through the UCSC Cell Browser from the “Human Cortical Development” collection (https://pre-postnatal-cortex.cells.ucsc.edu). First, clusters annotated as non-neuronal (“INT” and “Progenitors”) were removed, resulting in a dataset comprising 46,187 cells and 17,663 genes. Our neuronal clusters (C9, C10, C12, C14, C15, and C21, total cells = 2,012) were then integrated with the remaining interneuron clusters using Seurat-CCA Integration, after scaling and PCA reduction using default parameters (Seurat v5.3.0). The resulting dataset, containing 48,199 cells and 14,182 overlapping genes, was clustered using the Seurat pipeline with 19 principal components as input dimensions and a clustering resolution of 2.

To compare gene expression patterns between our most developed SST and PVALB neurons and Velmeshev *in vivo* counterparts, SST and PVALB cells were identified and extracted from the Velmeshev dataset by selecting cells belonging to the following annotated clusters: SST (“SST”, “SST_RELN”) and PVALB (“PV”, “PV_MP”). This subset comprised 23,806 cells and 17,663 genes. Differential gene expression between SST and PVALB cells was assessed using the FindMarkers function in Seurat (v5.3.0). Genes were considered significantly differentially expressed if they showed a fold change >2 and a Bonferroni-corrected p-value <0.05. Fisher’s exact test was used to evaluate the overlap between *in vitro* and *in vivo* DEGs for SST and PVALB cells relative to a background set comprising genes present in both the *in vitro* and *in vivo* datasets (6,175 genes). Statistical significance was defined as a Bonferroni-corrected p-value <0.05.

### Functional over-representation test (gene set overlap)

The degree of overlap between pairs of gene sets was evaluated using Fisher’s Exact test, where the background set consisted of all tested protein-coding genes (n=17,220).

### Statistical analysis and data presentation

Unless specifically stated in each methodology section, GraphPad Prism (version 10.6.1) was used to test the statistical significance of the data and to produce the graphs. Either One-way or Two-way ANOVA tests were used with Bonferroni-corrected post hoc tests (see figure legends for further test results). All data are presented as mean ± SEM with individual biological repeats as a dot.

## Acknowledgments

This work was supported by MRC Centre grant (MR/L010305/1), Waterloo Foundation ‘Changing Minds’ programme, the Royal Embassy of Saudi Arabia in the UK, and the start-up funding from the Neuroscience and Mental Health Research Institute, Cardiff University. We acknowledge excellent technical support for the next-generation sequencing from Joanne Morgan (MRC Centre, Cardiff University) and single-cell transcriptome library preparation from Angela Marchbank (Genome Hub, Cardiff University). We appreciate excellent general lab support from Emma Dalton, Trudy Workman and Olena Petter. We thank Prof. Meng Li and Dr. Claudia Tamburini for their advice and technical support in the initial stages of the project and Prof. Jeremy Hall for his support throughout the project. The graphical abstract was partially created in BioRender. Shin, J. (2026) https://BioRender.com/g55twtj

## Author contributions

**KA**: investigation, formal analysis, funding acquisition, visualisation, writing – original draft preparation, writing – review & editing **DDA**: data Curation, Formal Analysis, visualisation, writing – original draft preparation, writing – review & editing **AG**: investigation, formal analysis, funding acquisition, writing – review & editing **SW**: investigation, formal analysis, **AJP**: supervision, visualisation, writing – original draft preparation, writing – review & editing, **ES**: conceptualisation, formal analysis, funding acquisition, supervision, visualisation, writing – original draft preparation, writing – review & editing

## Declaration of interests

The authors declare no competing interests.

---

**Figure S1.**
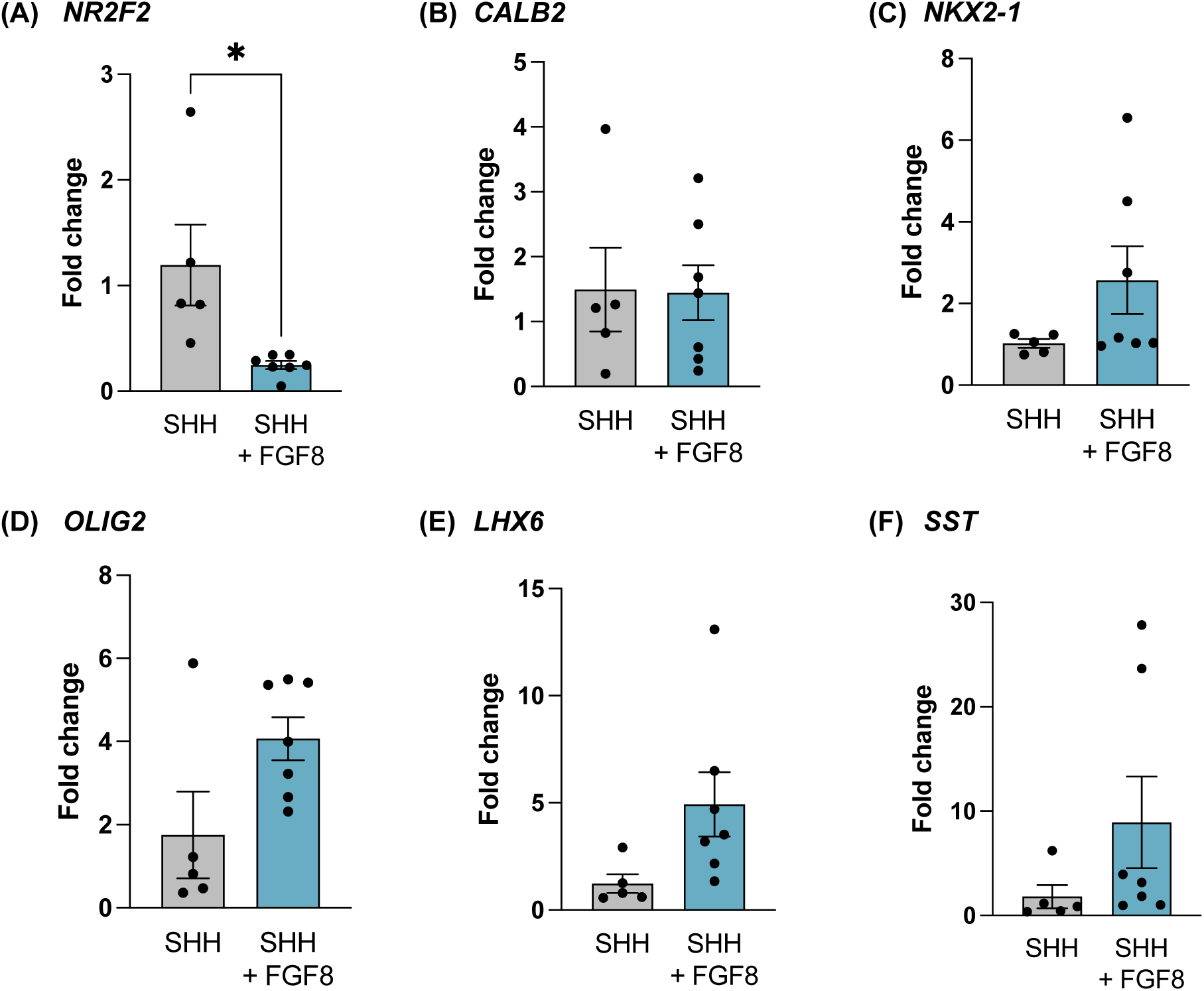
FGF8 rostralises cells, increasing the MGE-fated cells at the expense of the CGE-fated. H7 hESCs were differentiated with/without FGF8 from day 10 to 19 and harvested on day 50, and qRT-PCR was performed. CGE-expressed genes *NR2F2* (*COUTFII*, **A**) showed significantly decreased expression when treated with FGF8, although no difference was shown in *CALB2* (*CALETININ*, **B**) expression. Meanwhile, MGE-expressed genes showed an increased trend of expression - *NKX2-1*, (**C**), *OLIG2* (**D**), *LHX6* (**E**) and *SST* (**F**).

**Figure S2.**
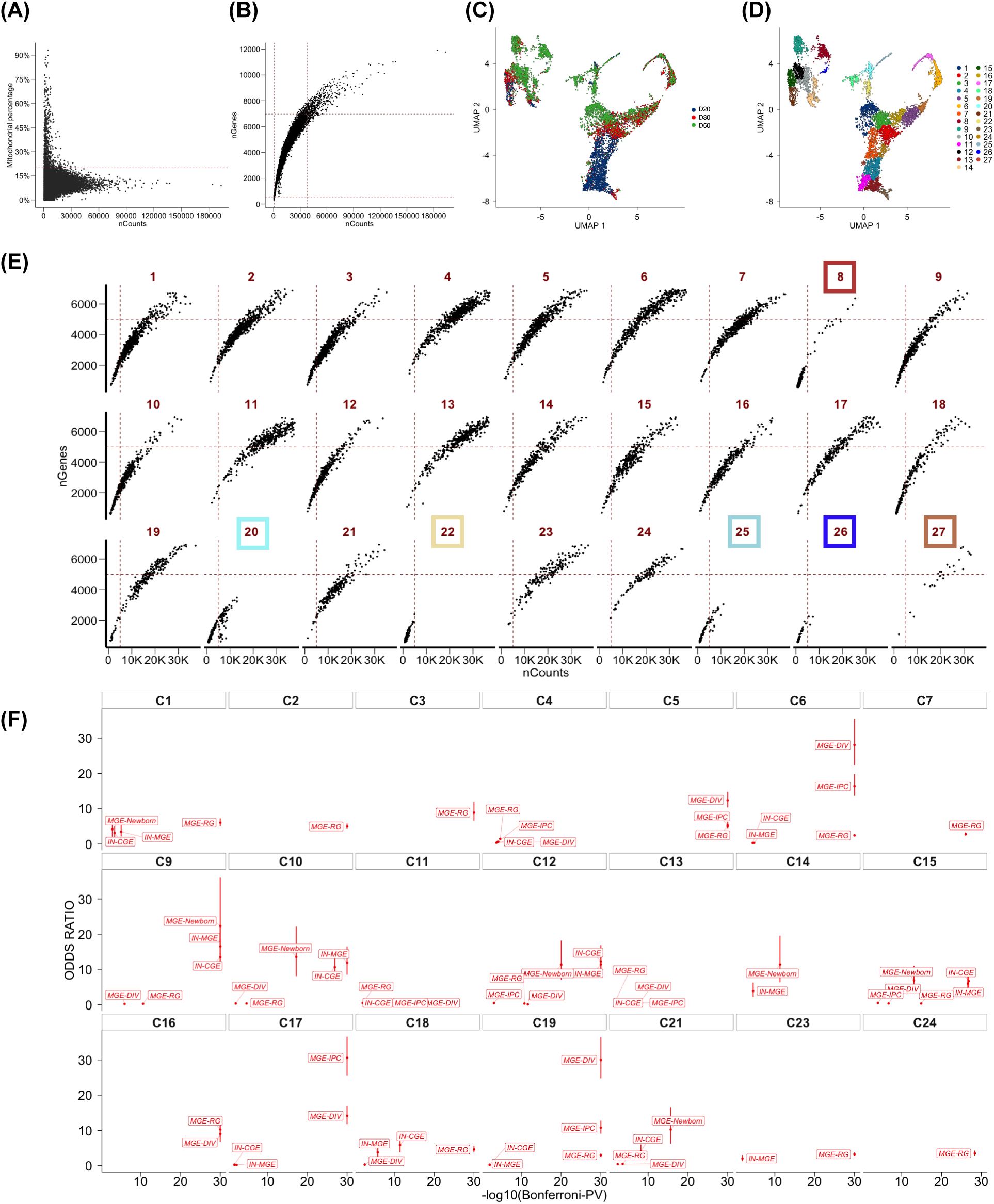
Cell and cluster quality check and cluster identity analysis. **(A)** Plots showing the number of counts versus the percentage of mitochondria genes. The threshold was set as 20%. **(B)** Plots showing the number of counts versus the number of genes after the mitochondria gene filteration. Cells that had more than 5% but less than 95%were included to filter out dead cells and doublets. **(C)** Clusters with *in vitro* differentiation days. **(D)** UMAP shows 27 cluesters. **(E)** 6 cluesters that were consisted of low quality (less number of counts and genes) were removed from the further analysis. **(F)** Identity of clusters was defined by enrichment analysis between DEGs from Nowakowski dataset and DEGs from our clusters. Odds Ratios and -log10(Bonferroni-p-value) for each cluster were plotted. The lines show the lower and higher odds ratio values. Only enrichments with Bonferroni p<0.05 are shown.

**Figure S3.**
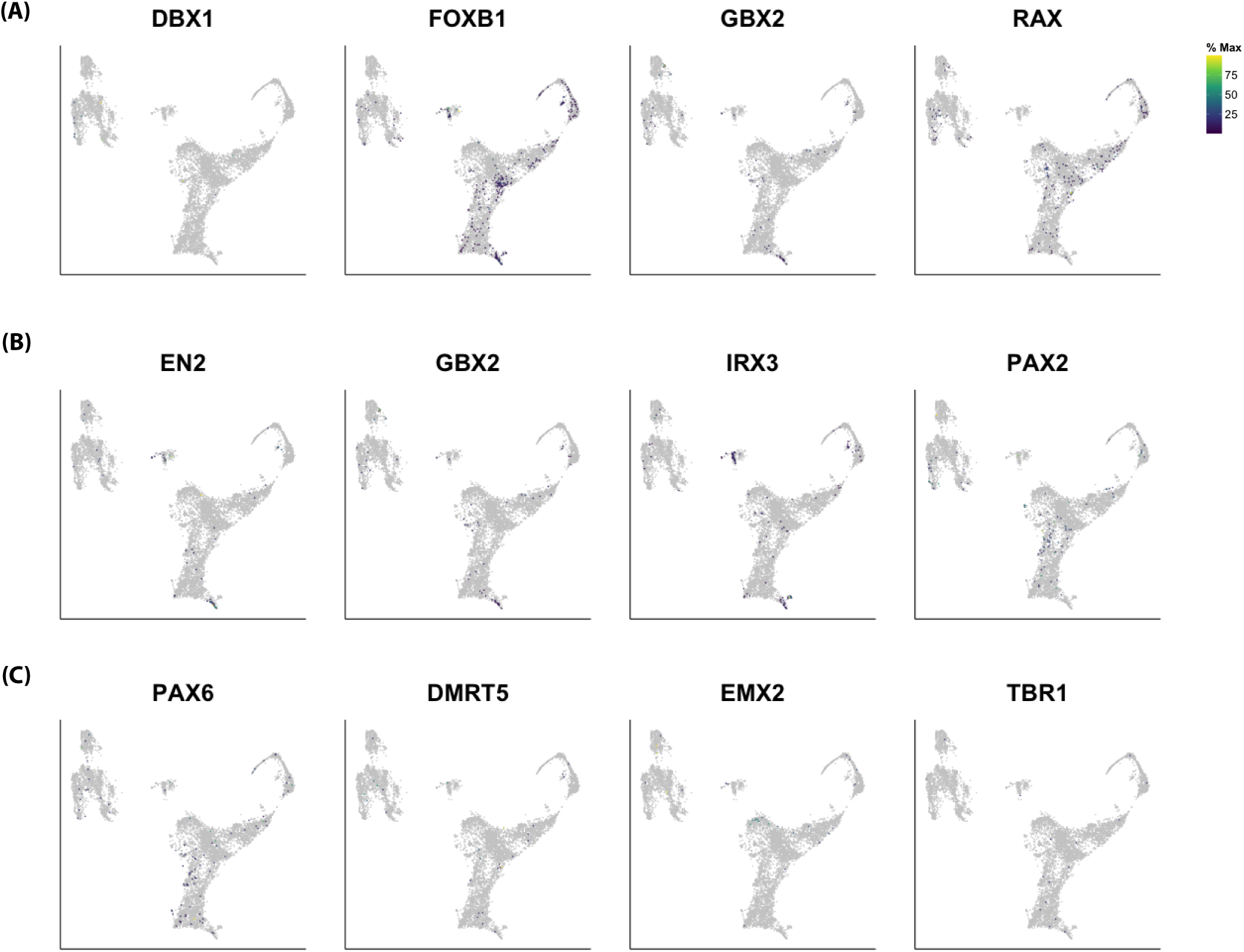
Canonical marker gene expression. **(A)** Diencephalic gene expression. **(B)** Midbrain and hindbrain gene expression. **(C)** Dorsal telencephalic gene expression.

## Notes

### Competing Interest Statement

The authors have declared no competing interest.

